# SURFBAT: a surrogate family-based association test building on large imputation reference panels

**DOI:** 10.1101/2023.01.10.523404

**Authors:** AF Herzig, S Rubinacci, G Marenne, H Perdry, FrEx Consortium, FranceGenRef Consortium, C Dina, R Redon, O Delaneau, E Génin

## Abstract

Genotype-phenotype association tests are typically adjusted for population stratification using principal components that are estimated genome-wide. This lacks resolution when analysing populations with fine structure and/or individuals with fine levels of admixture. This can affect power and precision, and is a particularly relevant consideration when control individuals are recruited using geographic selection criteria. Such is the case in France where we have recently created reference panels of individuals anchored to different geographic regions. To make correct comparisons against case groups, who would likely be gathered from large urban areas, new methods are needed.

We present SURFBAT (a SURrogate Family Based Association Test) which performs an approximation of the transmission-disequilibrium test. Our method hinges on the application of genotype imputation algorithms to match similar haplotypes between the case and control groups. This permits us to approximate local ancestry informed posterior probabilities of un-transmitted parental alleles of each case individual. SURFBAT provides an association test that is inherently robust to fine-scale population stratification and opens up the possibility of efficiently using large imputation reference panels as control groups for association testing. The method is suitable when the control panel spans the local ancestry spectrum of the case-group population and each control has similar paternal and maternal ancestries. This is the case for our reference panels where individuals have their four grand-parents born in the same geographic area. In contrast to other methods for association testing that incorporate local-ancestry inference, SURFBAT does not require a set of ancestry groups to be defined, nor for local ancestry to be explicitly estimated.

We demonstrate the interest of our tool on simulated datasets created from the 1000 Genomes project and the FranceGenRef project, as well as on a real-data example for a group of case individuals affected by Brugada syndrome.

## Introduction

Genome-wide association studies (GWAS) have become the established approach for an agnostic search of genes or genetic variants that may play a role in the development of complex multifactorial diseases [1]. A widely accepted notion is that in a case-control design, adjustments should be made for population stratification [2]. Confounding due to population stratification is avoided when using family-based association tests [3] where case individuals and their unaffected close relatives are both recruited to the study; hence each case individual will essentially have ancestry-matched controls. However, family-based designs are currently not widely used in the study of complex trait genetics due to the difficulty of recruiting large numbers of families. Indeed, due to the cost and infrastructure required for genome sequencing, a prevalent approach is to recruit only cases and compare them to external panels of controls or to set-up large cross-sectional studies (or biobanks) which allow for the study of many different phenotypes. There are many examples of population-based panels that could allow for association studies to be completed without the need to recruit further control individuals; e.g. the UK biobank [4], the Estonian biobank [5], goNL in the Netherlands [6].

The huge sample sizes that can be achieved with case-control or biobank designs give such power for the detection of new signals is the main reason that they have become far more prevalent that family based designs; but at the cost of having to hence deal with population stratification. This is usually achieved by adjusting for principal components calculated from a genotype correlation matrix or for fine-structure cluster membership. Such adjustments are ‘global’ in the sense that the principal components added to the association model use genome-wide calculations. It is possible to adjust locally, with the idea that patterns of stratification may not be equal in all genomic regions. This approach has been shown to be of interest [7,8], notably in the study of admixed populations [9,10].

Typically, such local adjustment requires local-ancestry inference to be first performed using either hidden Markov modelling (e.g. HAPMIX [11], LAMP [12], or FLARE [13]) or recent methods that use random forests (RFMix [14]), dynamic optimization (Loter [15]), or even neural networks (LAI-net [16]). Such methods essentially colour or paint [17] each study individual’s haplotypes based on the similarity of haplotype segments with haplotypes in a reference set of individuals from different ancestral groups (that have to be defined at some point). The local ancestry colouring can then be incorporated into association testing, for example through logistic regression (Tractor [18]), mixed-modelling (asaMap [19]), or through joint testing of genotype and ancestry associations [20]. Including such information has been demonstrated to enhance association studies in terms of power for discovery [18,21], fine-mapping [22,23], and even for studies of interaction effects [24]. A key problem in these methods is the pre-defined choice of ancestry groups and estimation of local ancestry. For studies of recently admixed populations, this may be practical [25–30] but in studies of populations with fine-scale population structure it will not be clear how to define different ancestry groups. For example, the French population harbours important fine-scale population structure [31] which should be taken into account during association testing. Yet it would not be clear how to best divide reference individuals into different groups, hence making local-ancestry inference problematic; a more fluid method is required.

Here we present a new method providing the following key advantages: our method adjusts for local patterns of population stratification, but unlike existing methods that do so, there is no constraint on having to choose and define ancestry groups or explicitly map local ancestry. This is achieved by approximating a family-based study design from case-control data using the idea of surrogate parents [32] enabling huge panels of control individuals to be exploited efficiently. This is accomplished by using Hidden Markov Models [33] (HMMs) that have been previously optimized in the domain of genotype imputation [34]. We describe our method as a SURrogate Family-Based Association Test (SURFBAT).

Genotype imputation methods are based on the Li-Stephens model [35] where given a large group of nhaplotypes in a population, an *n* + 1^*th*^ haplotype can be modelled as a mosaic of small haplotype chunks from the pool of *n*. In isolated populations (such as Iceland), this works particularly well due to longer sharing of haplotype segments that are identical-by-descent (IBD) [36,37]. The concept of surrogate parents suggests that for each given individual at a given point of the genome, even if the true parents of the individual are not present in the sample, two groups of surrogate parents can be identified who share a short haplotype that is at least very similar to the maternal or paternal haplotype of the given individual. This was originally employed for the purposed of statistical phasing [38], genotype imputation [39] and parent-of-origin analyses [40]. Similar ideas have recently resurfaced for the same themes [41– 43] now that biobank size data have become large enough that such approaches that were previously only viable in the domain of isolated populations have become applicable also for non-isolated populations.

SURFBAT takes a group of case individuals and identifies surrogate parents from within a large panel of control individuals. Crucially, we interrogate the haplotypes of the surrogates that are not shared with the case individuals in question. The non-shared haplotypes between case individuals and their surrogates, we argue, represent a resource of ancestry matched haplotypes and in the surrogate-parent interpretation represent an approximation of the un-transmitted alleles of the case individuals’ parents. This effectively provides a rough imputation of parental genotypes and hence an approximation of a Transmission-Disequilibrium Test (TDT) [44] can be made to test for association with a trait without sequencing parental genomes. Another interpretation of the method is that for each case individual, SURFBAT creates a pseudo-control matched on local-ancestry from within the control panel. SURFBAT performs a TDT test based on the methods that incorporate genotype uncertainty (as we are using imputed genotypes) given by Taub et al, [45]. SURFBAT locates and weights the contribution of surrogates using the Li-Stephens model for genotype imputation that is used by leading software IMPUTE5 [46]. A fuller description of the calculations of SURFBAT are given in the methods.

We demonstrate the properties of SURFBAT through simulation using the 1000 Genomes project [47] (1000G) and data from 856 individuals with Whole Genome Sequencing (WGS) data from the FranceGenRef project [48,49] (FGR). This is followed by a demonstration using true data from 346 case individuals diagnosed with Brugada syndrome [50].

## Results

For both the simulation study and real-data example, we will first define a control group. In both analyses our panel of control individuals involves 2504+856 individuals or 5008+1716 haplotypes attained by merging the 2504 individuals from the 1000G project with 856 individuals with WGS data from FGR. The steps used for construction of this panel are given in the Methods. The final dataset involves 10,252,495 bi-allelic variants across the 22 autosomal chromosomes.

### Simulation Study

We compared SURFBAT against a traditional GWAS adjusted on six principal components (Figure 1) using a simulation set-up devised to demonstrate the properties of our method. To this end, we constructed a group of 450 case individuals as mosaics of the control group. Mosaic construction was performed with R-package Mozza (https://github.com/genostats/Mozza) and is described in the Methods. For the purposes of the simulation, we only simulated a short genome (chromosomes 10-22). The case individuals have an admixed ancestry profile with 50% of their chromosome chunks coming from the AFR populations of 1000G (African continent) and the other 50% from the French samples of FGR. We added a signal of purely local ancestry on Chromosome 11 by simulating an excess of FGR haplotypes, and a more specific signal on Chromosome 20, with an excess of FGR haplotypes carrying the alternative allele for the variant rs197819 (chosen randomly).

**Figure 1:**
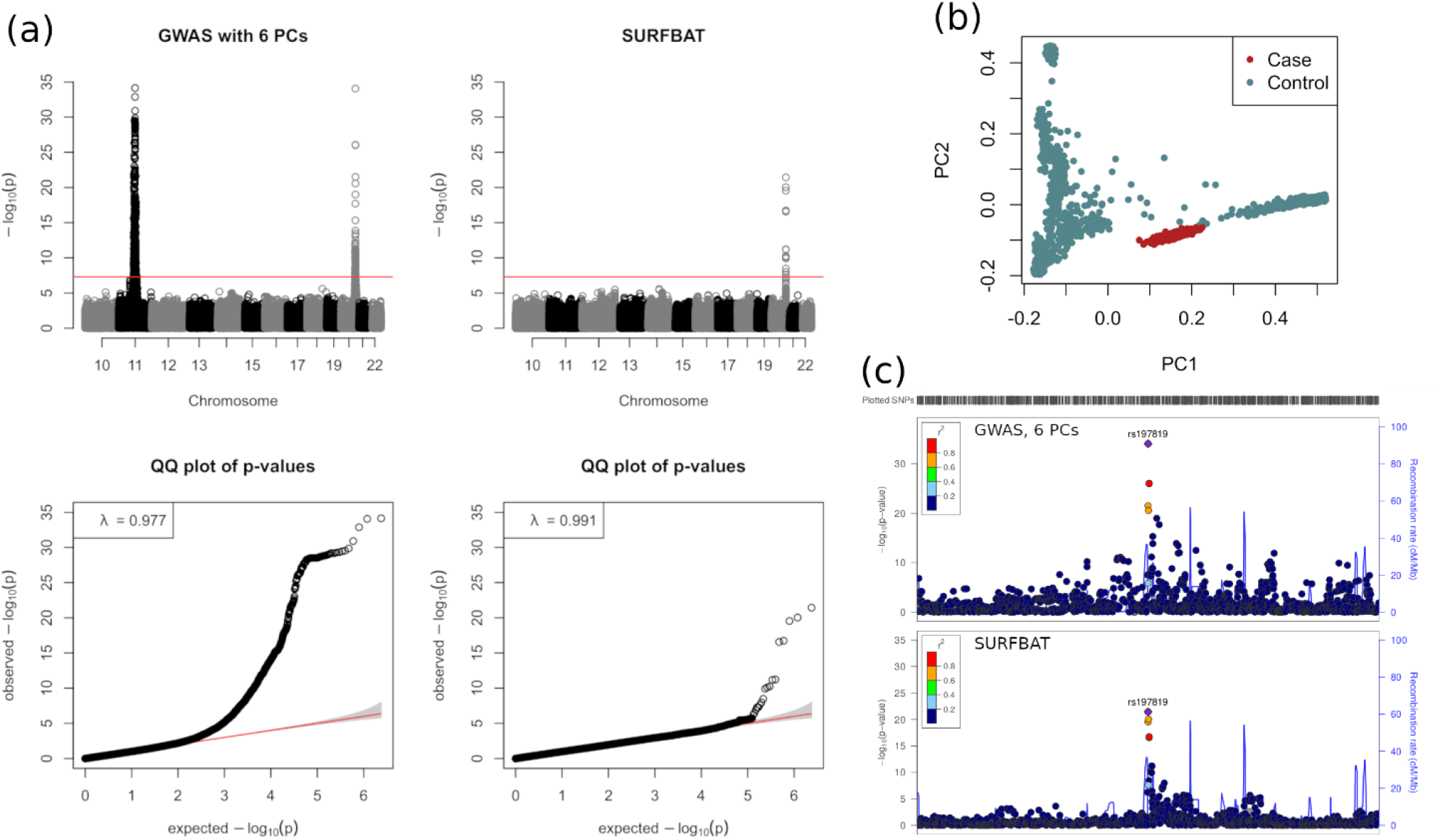
(a) Manhattan and QQ-plots of association studies either carried out using a standard GWAS (logistic regression) adjusted for six principal components (left plots), or with SURFBAT (right plots). Red lines indicate genome wide significance (5e-8). (b) Principal components 1 and 2 estimated on the all simulated cases (red) and controls (blue) (individuals of the 1000 Genomes project and FranceGenRef). (c) LocusZoom [51] of the signal on chromosome 20 for the two methods.

Here we observe the two key properties of SURFBAT, local ancestry is adjusted for and hence the strong GWAS signal on Chromosome 11 disappears. The association signal on Chromosome 20 is detected and is also noticeably more precise as we deliberately combined the association with rs197819 with an association with local ancestry (see Methods). Indeed, with a direct GWAS, there are SNPs that surpass genome-wide significance up to 0.5Mb away from rs197819; whereas SURFBAT only gives significant p-values to variants close to rs197819 (Figure 1c).

### Analysis on real data with cases affected by Brugada syndrome

We examined an example of real data using 346 cases with Brugada syndrome. These individuals were recruited in France and comprise part of the case group involved in the largest GWAS to date for Brugada syndrome [50]. Three different possible scenarios of an association study were tested. (i) We performed a GWAS, adjusted on principal components (six to match the GWAS in Barc et al., [50]), against the aforementioned control group of 1000G+FGR. (ii) We applied SURFBAT, using 1000G+FGR as the control group and only array data for the 346 Brugada-case individuals. (iii) A GWAS of the 346 case individuals against 569 French control from the FrEX cohort [52,53] (http://lysine.univ-brest.fr/FrExAC/). In this third scenario, both the Brugada-case group and FrEx control group had only array data and were both imputed using 1000G+FGR as an imputation panel. In Figure 2, the results of these three strategies are shown, and in Supplementary Table 1 the p-values for leading SNPs from the GWAS of Barc et al., [50] are given and compared against our results. As we have a far smaller sample size (Barc et al., [50] analysed 2,820 cases and 10,001 controls compared to our 346 cases and 3360 controls), we did not have the power to detect signals aside from the two most significant at SCN10A on chromosome 3 and HEY2 on chromosome 6. As shown in Supplementary Table 1, the association analyses that we performed, including SURFBAT, were however able to replicate (p<0.05) other GWAS hits from Barc et al, [50]. Genomic control [54] was applied to all three association studies as SURFBAT’s test statistics (in particular) were inflated in this analysis. Indeed, SURFBAT may sometimes require genomic control, a full discussion of this is given in the Methods.

**Figure 2:**
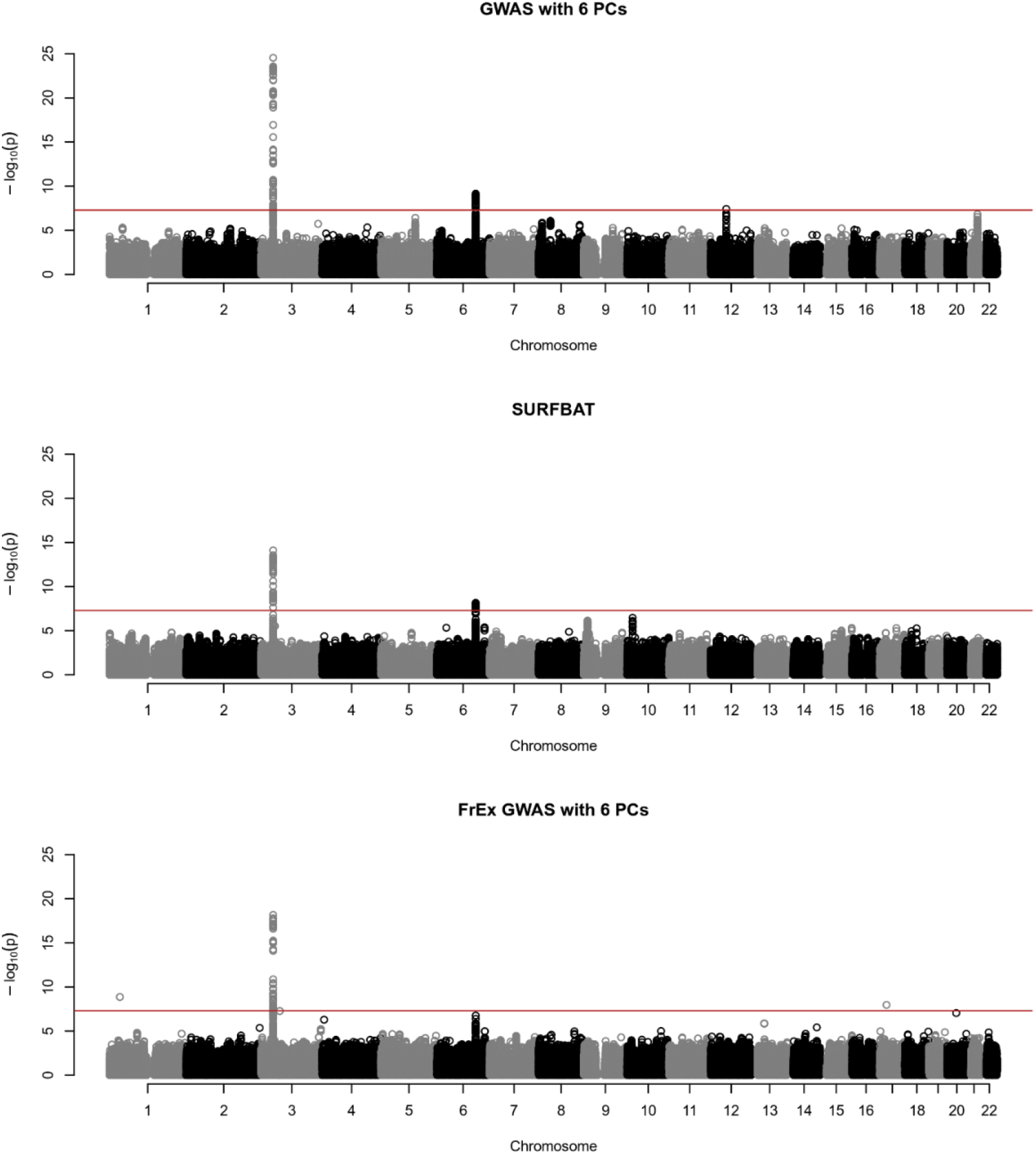
Association analyses of the Brugada dataset. Top: genome-wide association study (GWAS) comparing 346 Brugada-case individuals with 3360 control individuals (FGR+1000G) adjusted for six principal components (PCs); both cases and controls have whole-genome sequencing (WGS) data. Middle: SURFBAT, 346 Brugada-case individuals with only genotyping array data are compared to the WGS data control group (FGR+1000G). Bottom: GWAS of 346 Brugada-case individuals and 569 FrEx-controls individuals; here both cases and control have genotyping array data only and both are imputed using WGS data of the control group FGR+1000G; here we adjusted for six PCs calculated on the genotyping array data. All association analyses results are given for the same set of 5,462,920 variants; having excluded variants with a minor-allele frequency below 1% in the case group or with an imputation ‘info’ score below 0.4. Red lines indicate genome wide significance (5e-8).

## Discussion

Here we have presented a novel method, SURFBAT, for association testing for common genetic variants with binary phenotypes. SURFBAT compares a group of case individuals against a large reference panel of control individuals using the hidden Markov modelling that have been demonstrated to be highly effective for genotype imputation. This is achieved by estimating allelic dosages for the un-transmitted alleles of each case individual; hence with a test statistic derived from the literature of family based studies. This allows SURFBAT to adjust for population stratification by design, and not just at a global level but down to a local ancestry level and hence should be an interesting tool for the study of admixed individuals. This is shown in the simulation study where a large GWAS peak (chromosome 11) coming only from local ancestry completely disappears using SURFBAT whilst a ‘true’ peak (chromosome 20) is maintained. Importantly, local ancestry in taken into account without having to specify a number of ancestry groups or to map local ancestry. Furthermore, the signal on chromosome 20 was much cleaner than in the GWAS suggesting that SURFBAT could aid in studies of fine-mapping.

SURFBAT takes advantage of the highly efficient software IMPUTE5, hence this could allow for huge imputation reference panels (such as the HRC or even TOPMED) to be used as control panels for GWAS. As IMPUTE5 will search out the most appropriate haplotypes in the reference panel for each target haplotype; the presence of haplotypes in the reference panel that are not highly relevant to the imputation of a given target haplotype is not problematic. In this aspect, SURFBAT therefore resembles the UNICORN method proposed by Bodea et al, [55] where case individuals are compared to a large ensemble of reference individuals. In both the simulation study and real data example presented here, performing a traditional GWAS adjusted for principal components was in fact more powerful than SURFBAT. However, in many cases such a strategy of directly performing a GWAS would not be possible and/or tractable as individual-level data for the largest imputation panels are not publically available. However, SURFBAT could easily be performed at the same time as imputation opening up the possibility for online imputation servers such as the one in Michigan [56] to also facilitate GWAS without having to share their data. This is demonstrated in the analysis of the Brugada dataset where SURFBAT is able to achieve similar results to a direct GWAS against the control group of 1000G+FGR but without the requirement of jointly manipulating the case and control data together in order to calculate PCs and to perform the association testing. Even if individual-level data are available, for a huge control panel, merging the data with the case group, calculating PCs, and performing the GWAS can all be highly time consuming.

Another key advantage of SURFBAT is that it only requires genotyping array data for case individuals, hence allowing for a very simple case-only designs where instead of having to spread a sequencing budget across cases and controls, one could simply genotype a large number of case individuals and compare them to an appropriate existing imputation reference panel using SURFBAT. This could also prove advantageous when grouping case individuals together from many different cohorts. Indeed, when comparing SURFBAT to the FrEx GWAS study design, where both cases (Brugada) and controls (FrEx) are imputed, SURFBAT achieved genome-wide significance for the signal on HEY2/NCOA7 whereas the FrEx GWAS did not.

A further potential application would be that SURFBAT could also be used to ‘complete’ families, as in [57,58] where methods were presented for including singleton cases in family-based designs using simple imputation methods. SURFBAT could essentially be used for the same approach but with a more elaborate method for estimating both transmitted and un-transmitted allelic dosages.

SURFBAT therefore provides an interesting counterpoint to GWAS and is very practical to put in place as we have constructed the test so that it can be achieved simultaneously to the imputation of genotypes. As imputation reference panels have become incredibly large, SURFBAT provides a methodology to harness such panels for association testing efficiently. There are however, certain limitations to SURFBAT. As noted, SURFBAT was less powerful than GWAS in both the simulation and real data example. This diminished power results from the fact that we are essentially comparing K cases with K pseudo-controls but also because we are performing a paired test. A gain in power was however observed if we perform a simple unpaired test (see Methods and Supplementary Figure 1) and SURFBAT is equipped with optional functionality to perform this unpaired test. Furthermore, here we have developed a test for binary traits under an additive genetic model for common genetic variants. Though, it would be straight forward to extend the method for quantitative traits, other genetic models, adjusting for covariates, and even to perform burden tests for association with rare variants. It must also be mentioned that the performance of SURFBAT depends entirely on the availability of an appropriate imputation reference panel for one’s set of case individuals. The composition of the imputation reference panel is a very important consideration for genotype imputation [59] and will therefore also be important for SURFBAT. Finally, we saw in the real data example that SURFBAT could return inflated test statistics; this was dealt with using genomic control. This inflation was however not apparent on the simulated dataset; certainly because in this instance the case individuals were simulated as mosaics of the control individuals and hence the surrogate parent model was very effective. As imputation reference panels increase in size, surrogate parents should be more readily found and hence SURFBAT should not suffer from inflation.

## Methods

### The SURFBAT approach

In standard genotype imputation, each haplotype (*h*_*i*_) of the target group is modelled as an imperfect mosaic of the 2*N* haplotypes (*H*_*k*_, *k* = 1, . . ., 2*N*) of the reference panel. This is achieved using an HMM, with states at each genomic position where a variant is observed in both the reference panel and the target group; so typically the list of positions for which the target group have been genotyped. The hidden states of the model at position *j* indicate which haplotype (or cluster of haplotypes) in the reference panel is providing the mosaic tile for haplotype *h*_*i*_ at position *j*; often referred to as the copying state as the target haplotype will probabilistically ‘copy’ from these reference haplotypes to achieve the missing genotype imputation. The observed states are the allelic values of haplotype *i*. The implementation of the HMM and the transmission and emission probabilities can slightly vary between software; globally the transition probabilities are calculated based on the size of the reference panel and an estimated recombination rate between the genetic locations of adjacent states; and the emission probabilities allow for the mosaic to be imperfect in the sense that the target haplotype and the haplotypes in the reference panel can differ due to either recent mutations or genotyping error.

We denote the hidden states of haplotype *h*_*i*_ as *s*_*i*_ which take values in 1, …, 2*N* where *N* is the number of diploid individuals in the reference panel. The observed states (allelic values coming from genotyping data) of haplotype *h*_*i*_ are denoted as *o*_*i*_. The HMM provides the posterior probabilities of the hidden states at each position *j* using the forward-backward algorithm [33,60]: 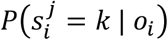. These probabilities are then extended to markers, noted as *j*′, that are not shared between the target and reference panel through linear interpolation.

The missing alleles for marker *j*′ in haplotype *i* are then imputed with the following dosage:

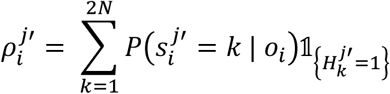

Where the quantity 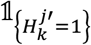 is equal to 0 or 1 if the reference haplotype *H*_*k*_ carries a major or minor allele at marker *j*′, respectively. The final imputed dosage for individuals in the target group is simply the sum of the dosage of their two haplotypes. This dosage will take a value between 0 and 2 and represents the individual’s expected count of minor alleles for marker *j*′. However, here we are interested in the haplotype dosages. SURFBAT will keep each individual’s two haplotype dosages separate and furthermore will calculate what we will term the un-transmitted dosages which are simply as follows:

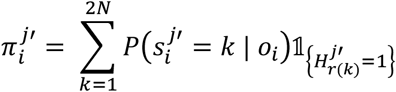

Assuming that the haplotypes in the reference panel are stored in groups of two for each reference panel individuals (reference haplotypes 1 and 2 correspond to the first individual, haplotypes 3 and 4 to the second individual etc.), *r*(*k*) is a simple function that gives the index of the partner haplotype to haplotype *k*. Explicitly, *r*(*k*) = *k* + 1 if *k* is odd and *r*(*k*) = *k* − 1 if *k* is even. We implemented the calculation of both 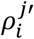 and 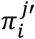 within the existing imputation algorithm IMPUTE5, for both positions that are not genotyped (e.g. *j*′) and that are genotypes (e.g. *j*), alike.

Note that there is an implicit assumption that the two haplotypes within an individual on the control panel share a relatively similar ancestry - in the context of our study, this is reasonable for the 1000G individuals and is also respected by the FranceGenRef panel by design [47–49]. This is because the individuals of FranceGenRef were recruited based on grand-parent birthplace data taking individuals with all four grand-parents born within a small locality hence approximately insuring that such individuals have both maternal and paternal haplotypes from a similar region. Indeed, such a recruitment strategy has often been used for the construction of other reference panels [61–63]. Therefore, when the reference panel has such a construction, haplotype *H*_*r*(*k*)_ will be approximately matched (in terms of ancestry) to haplotype *H*_*k*_.

When using IMPUTE5 with the specifically designed option ‘--surfbat’, then at marker *j*′ for target individual *i* (with haplotypes *i*_1_ and *i*_2_), the four dosages 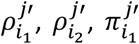 and 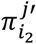 are calculated. In the ‘surrogate family’ interpretation of the four dosages, individual *i* has received two alleles with expected values 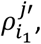, and 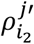 from their parents and the corresponding un-transmitted alleles from the parents have expected values 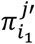 and 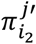. Then using similar notation to Taub et al, [45] we form the following test statistic for equilibrium of transmission at marker *j*′:

First, the quantities *NUM* and *DEN* are calculated:

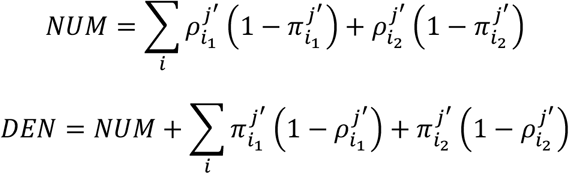

Then we calculate: 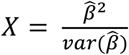 where 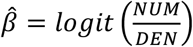, and 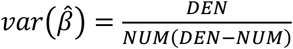. *X* will asymptotically follow a chi-squared distribution with 1 degree of freedom under the null hypothesis of equilibrium of transmission. This corresponds to an exact-form test statistic for conditional likelihood regression [64].

We also provide an alternative test where the cases and pseudo-controls are not paired; a simple test of marginal homogeneity [65] from a contingency table of the expected allelic values of the haplotypes of the case individuals and of the pseudo controls. The test is constructed from a contingency table as follows:

|  |  |  |
| --- | --- | --- |
| A | C | $n_{1.}$ |
| B | D | $n_{2.}$ |
| $n_{.1}$ | $n_{.2}$ | |

Where *n*_1_ = *A* + *C, n*_2_, = *B* + *D, n*_1_ = *A* + *C, n*_2_ = *C* + *D* and

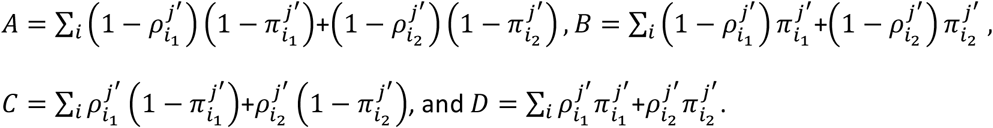

Then 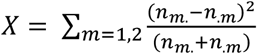 will again asymptotically follow a chi-squared distribution with 1 degree of freedom under the null hypothesis of marginal homogeneity. A comparison of the two tests are given in Supplementary Figure 1 for the Brugada dataset example; the unpaired test attributed smaller p-values to the SNPs that are likely not following the null hypothesis near SCN10A on chromosome 3 and HEY2 on chromosome 6.

In order to perform SURFBAT, only genotyping array data for the case individuals are required and WGS data for the controls individuals who must be formed into to a phased imputation reference panel. The cases are imputed against the controls using IMPUTE5 and the ‘--surfbat’ flag activated, which also calculates the per-SNP test-statistics and corresponding p-value.

There is the possibility to place thresholds on the minor allele frequency (MAF) and the imputation quality (INFO score), with default settings placed at 0.01 and 0.4 respectively. This is due to the fact that for rare variants and poorly imputed variants the test will not be appropriate; as is the case for a traditional GWAS using imputed data. We observed in the real data application presented in this work, an inflation of the SURFBAT p-values across the genome; similar to as observed in Taub et al, [45]. A demonstration of this is given in Supplementary Materials where the imputed dosage data from chromosome 3 for the 346 Brugada-case are compared against the dosages of their pseudo-controls (estimated by SURFBAT) using a principal component analyses (Supplementary Figure 2a). This inflation is likely due to the difference in the precision of imputation of the transmitted alleles (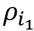, and 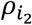) and the un-transmitted alleles 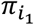, and 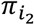). With a large enough reference panel, IMPUTE5 should be able to find a more precise mosaic of surrogate parents based on longer shared-haplotypes [42] and hence this difference in the precision demonstrated in Supplementary Figure 2a should be far less evident. The inflation can nonetheless be effectively adjusted for using genomic control [54] (Supplementary Figure 2b) and indeed was not observed in our simulation study.

### Creating a control panel

We took the haplotypes from the 1000G (Phase 3) from all populations, as were made available by imputation software IMPUTE2 [66]. This dataset was merged with the whole genome sequencing positions of FGR. In order to prepare the FGR data, we used the sequencing data quality control pipeline RAVAQ [67], and phased the data using SHAPEIT4 [68]. Merging and manipulation of the data file was achieved using the R-package gaston [69], plink v1.9 [70], and pbwt [71]; variants with a difference in minor allele frequency greater than 0.1 between FGR and the non-Finnish European 1000G individuals were removed from the control panel as these were deemed to likely represent a batch effect between the sequencing data of FGR and the public 1000G data. This panel can then be used either directly as a control group, or as an imputation reference panel for the purpose of SURFBAT. This was the control group/imputation panel for both our simulation study and real data example.

### Simulation study

Our simulated study involved simulating a group of 450 individuals who were constructed as mosaics of the control group using the R-package Mozza (https://github.com/genostats/Mozza) which leveraged the Li-Stephens Markov process of forming new mosaic haplotypes. To keep calculation time short, we only simulated chromosomes 10-22 and each chromosome was simulated separately. At each instance when a mosaic tile was drawn for a new case individual, we assigned a 50% chance to copy from a haplotype from the African continent super population from 1000G (AFR) and 50% from FGR. A tile length of 10cM was used for the simulation. We simulated a pool of 20,000 haplotypes; our 450 simulated case individuals’ 900 haplotypes were sampled from this pool in order to create two association signals on chromosomes 11 and 20. On chromosome 11, we sampled an excess of haplotypes with tiles copied from FGR haplotypes overlapping base-pair position 66602100 (near the middle of the chromosome). On chromosome 20, we sampled an excess of FGR haplotypes that carry an alternative allele for the common variant rs197819. This variant was chosen at random among a set of candidate variants with at least a MAF of 0.2 in both AFR and FGR. Genotyping data for the case group was simulated by extracting positions from the UK biobank SNP-array (details here: https://biobank.ndph.ox.ac.uk/showcase/refer.cgi?id=149601).

### Brugada study

We took 346 individuals with whole-sequencing data affected with Brugada syndrome as a real-data example. These individuals were recruited in the West of France and so we can assume that they can be accurately imputed using a reference panel built of FGR and 1000G [48]. To perform GWAS, we analysed all common bi-allelic variants (minor allele frequency above 0.01) that were present in our control panel (1000G+FGR). GWAS was carried out in the R-package gaston, using logistic regression adjusted for six principal components (PCs); PCs were calculated from pruned data and this was also achieved using gaston. To apply SURFBAT to the Brugada case individuals, we require only genotyping data for these individuals. We simulated this scenario by extracting positions from the UK BioBank SNP-array from the WGS data for the 346 individuals; this array was designed to facilitate genotype imputation and hence would be a logical choice for future applications of SURFBAT. The data from the extracted array positions were phased with SHAPEIT4 and then supplied to IMPUTE5 with the required option (--surfbat) in order to calculate genome-wide SURFBAT p-values. Finally, to demonstrate the interest of the concept of using an imputation reference panel as a control panel, we simulated a scenario where we have both case and control individuals with genotyping array data. In this circumstance, both cases and controls were imputed and a GWAS was performed on the imputed dosages. For this we took control individuals from 569 individuals of the FrEx database who have genotyping data (Illumina OmniExpressExome array). Positions corresponding to this array were extracted from the 346 case individuals and we then phased (SHAPEIT4) and imputed (IMPUTE5) the 346 case and 569 control individuals together using 1000G+FGR as a reference panel. GWAS was then performed on the imputed dosages using SNPTEST [72,73]; again using logistic regression adjusted for six PCs. All three association studies were corrected for inflation using genomic control [54] as we observed values of λ above 1 (Supplementary Figure 3); particularly for SURFBAT (see also Supplementary Figure 2a-b).

### Software Availability

SURFBAT will be made available as a new functionality in the existing software IMPUTE5 (https://jmarchini.org/software/) at the moment that the next software update is released in early 2023.

## Data Availability

Data from the FranceGenRef panel will be submitted to the French Centralized Data Center of the France Medicine Genomic Plan that is under construction. Enquiries for the use of this data can be addressed to GENMED LABEX (http://www.genmed.fr/index.php/en/contact). Those wishing to access the Brugada data on a collaborative basis should contact Richard Redon and Christian Dina. Summary information for the FrEx dataset is available at http://lysine.univ-brest.fr/FrExAC/, those wishing to access the data on a collaborative basis should contact Emmanuelle Génin.

## Author Contributions

AFH and EG devised the concept of the study. The methodology was developed by AFH. The choice of analyses and investigations included were discussed and devised by all authors: GM and HP for the simulation study, SR, GM, HP and OD for details of the method implementation, CD and RR for the Brugada analysis, AFH and EG for all of the above. Analyses were performed by AFH. The method was implemented by AFH and SR. AFH wrote the manuscript, all authors participated in the editing process and all approved the final manuscript. Funding for the work was secured by CD, RR, OD, and EG. Data for the study was provided by CD, RR, EG, the FrEx consortium and the FranceGenRef consortium.

