## Supplementary Materials for "SURFBAT: a surrogate family-based association test building on large imputation reference panels"

**1**: Inserm, Univ Brest, EFS, UMR 1078, GGB, Brest, France

**2**: Department of computational Biology, University of Lausanne, Lausanne, Switzerland

**3**: Swiss Institute of Bioinformatics, Lausanne, Switzerland

**4**: CESP Inserm U1018, Université Paris-Saclay, Villejuif, France

**5**: LABEX GENMED, Centre National de Recherche en Génomique Humaine, Evry, Paris

**6**: Inserm, unité de recherche de línstitut du thorax UMR1087 UMR6291

**7**: Université de Nantes, Institut National de la Santé et de la Recherche Médicale : U1087, Centre National de la Recherche Scientifique : UMR6291

**8**: CHRU Brest, Brest, France

* Corresponding Author

**Supplementary Materials**

| SNP | Gene | Barc et al, 2022^1^ | GWAS with 6 PCs | SURFBAT | FrEx GWAS  with 6 PCs |
| --- | --- | --- | --- | --- | --- |
| rs7638909 | SCN5A | **2.79e-8** | **2.70e-4** | **0.0184** | **1.45e-3** |
| rs62241190 | SCN5A | **8.56e-14** | **1.86e-3** | **1.21e-3** | 0.0111 |
| rs7374540 | SCN5A | **3.56e-57** | **4.77e-7** | **2.50e-3** | **1.13e-4** |
| rs7433206 | SCN5A | **9.52e-24** | 0.277 | 0.329 | 0.180 |
| rs34760424 | SCN5A | **3.03e-23** | **7.42e-4** | 0.0794 | **4.90e-4** |
| rs41310232 | SCN5A | **1.19e-15** | **2.45e-5** | **1.31e-3** | 0.0126 |
| rs6782237 | SCN5A | **1.05e-47** | **3.25e-8** | **3.71e-5** | **1.82e-6** |
| rs6801957 | SCN10A | **1.30e-180** | **2.89e-25** | **5.22e-14** | **6.89e-19** |
| rs6913204 | HDDC2 | **1.3e-8** | 0.313 | 0.0810 | 0.218 |
| rs9398791 | HEY2,NCOA7 | **1.49e-39** | **3.79e-9** | **1.03e-8** | **1.45e-6** |
| rs11765936 | TBX20 | **4.3e-11** | **4.32e-3** | **3.84e-3** | **4.01e-3** |
| rs340398 | TBX20 | **1.76e-9** | 0.351 | 0.492 | 0.0349 |
| rs804281 | GATA4 | **1.22e-9** | **1.23e-3** | 0.660 | 0.71 |
| rs72671655 | ZFPM2 | **2.51e-13** | **1.32e-3** | **9.34e-3** | **8.01e-3** |
| rs72905083 | WT1 | **2.09e-9** | 0.0790 | **0.0340** | 0.244 |
| rs883079 | TBX5 | **1.59e-10** | **0.0171** | 0.552 | 0.0575 |
| rs11645463 | IRX3 | **1.27e-9** | 0.0747 | 0.410 | 0.203 |
| rs72622262 | CRNDE,IRX5 | **1.37e-11** | 0.0789 | **0.0219** | **0.0339** |
| rs12945884 | PRKCA | **3.31e-8** | **0.0230** | 0.326 | 0.145 |
| rs476348 | MAPRE2 | **2.64e-9** | **0.0189** | 0.830 | **1.66e-3** |
| rs133902 | MYO18B | **7.73e-9** | **3.02e-3** | 0.449 | **1.24e-3** |

Supplementary Table 1: P-values from the three association analyses presented in Figure 2 in the main text for the set of SNPs identified in Table 1 of Barc et al.,^1^. We highlight in gold the p-values below 5e-8 and in blue those below 0.05.

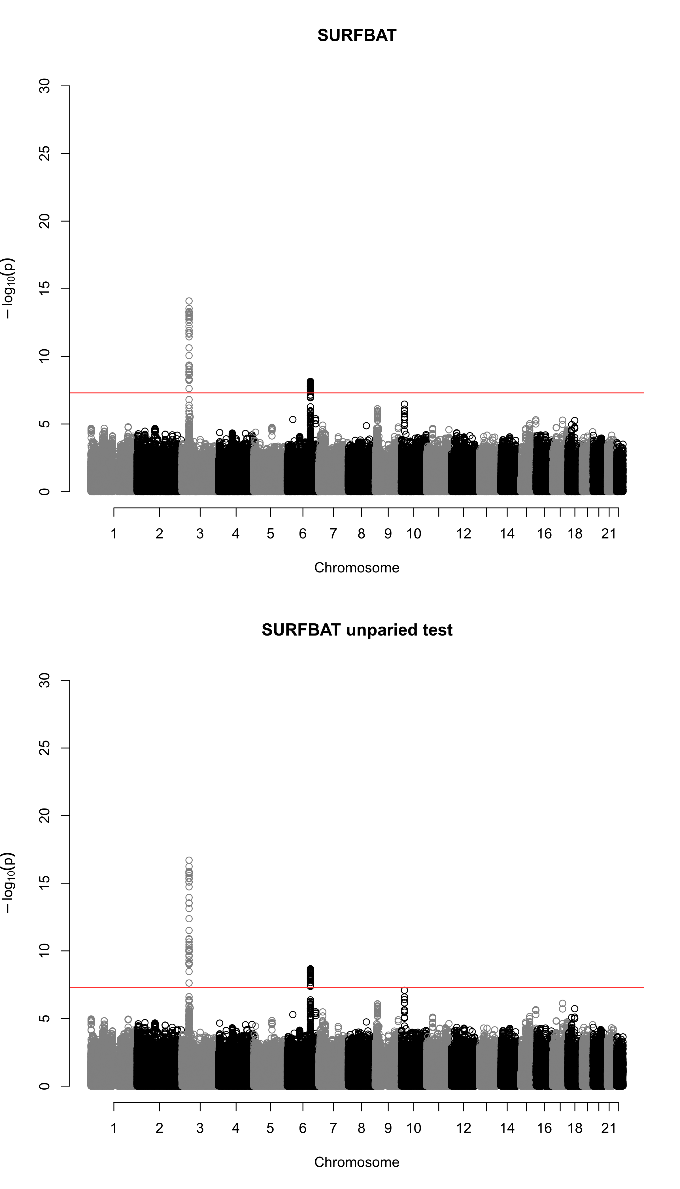

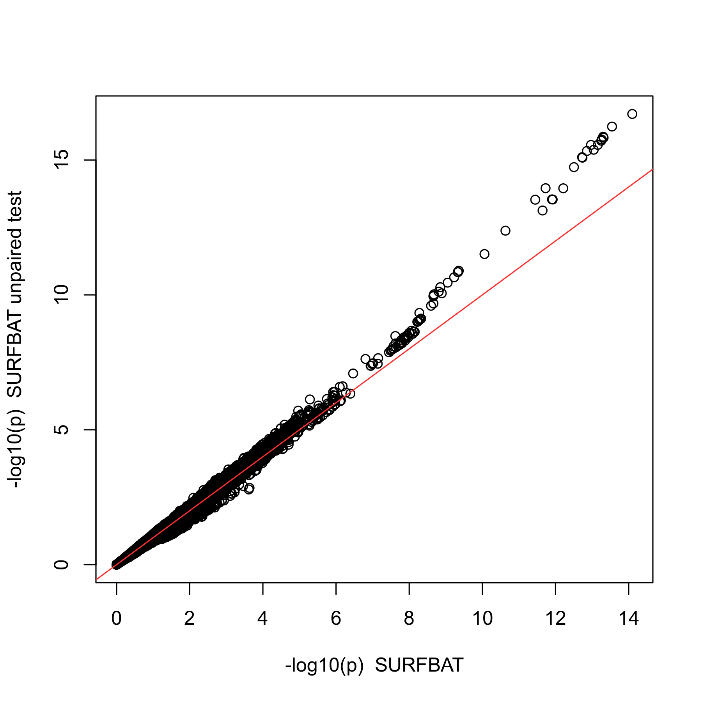

Supplementary Figure 1: Comparison of the SURFBAT with the unpaired version of the test. Genomic control has been applied in both sets of test statistics.

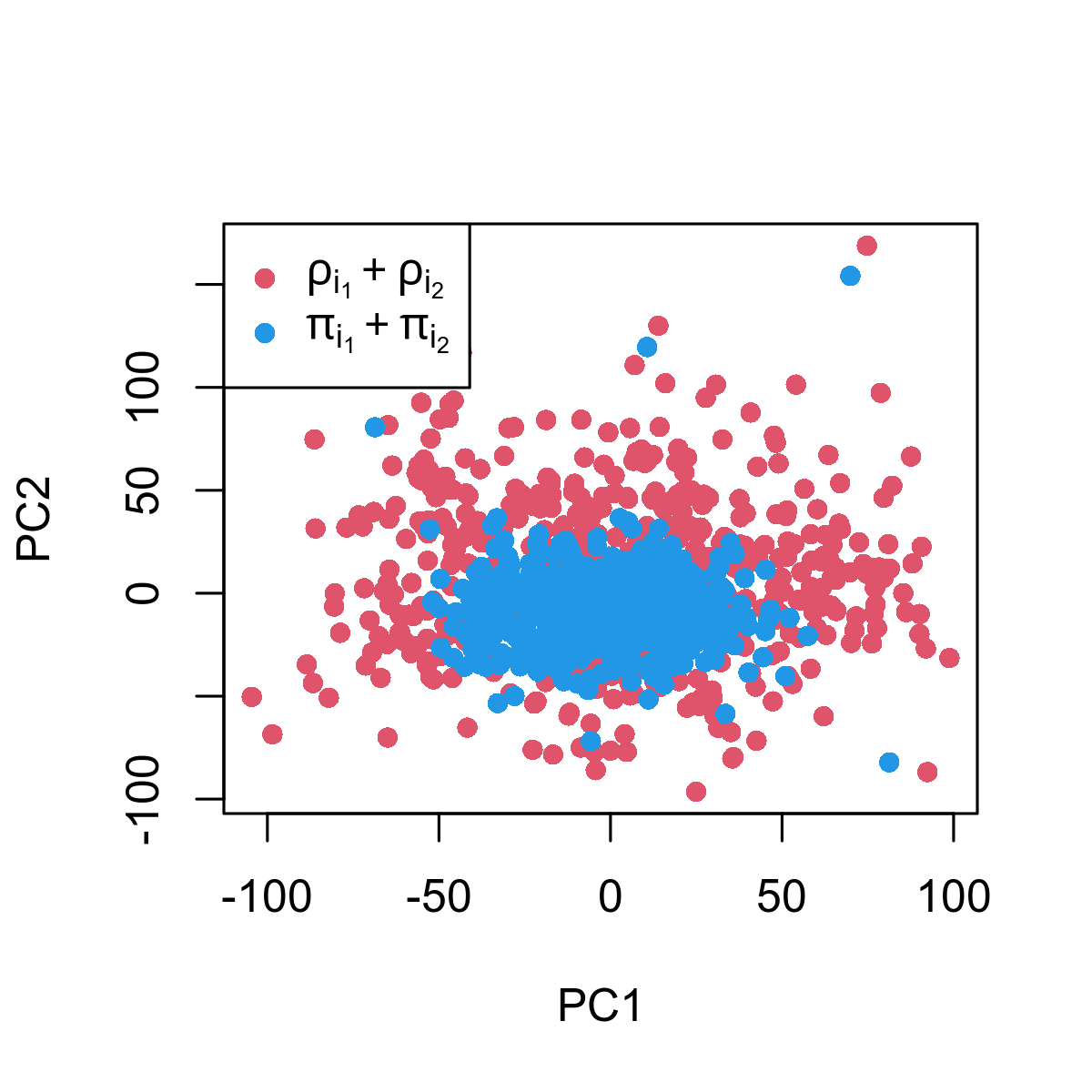

Supplementary Figure 2a: Principal components on chromosome 1 comparing the imputed dosages for the 346 Brugada-case individuals ($\rho_{i_{1}}$, +$\rho_{i_{2}}$) and the imputed dosages of their pseudo-controls using SURFBAT ($\pi_{i_{1}}$, and $\pi_{i_{2}}$). As the imputation of the un-transmitted alleles will naturally be less precise, we see less variability in this group; engendering a batch effect that could be compared to a comparison of imputed data from two different sources (eg. from different genotyping arrays).

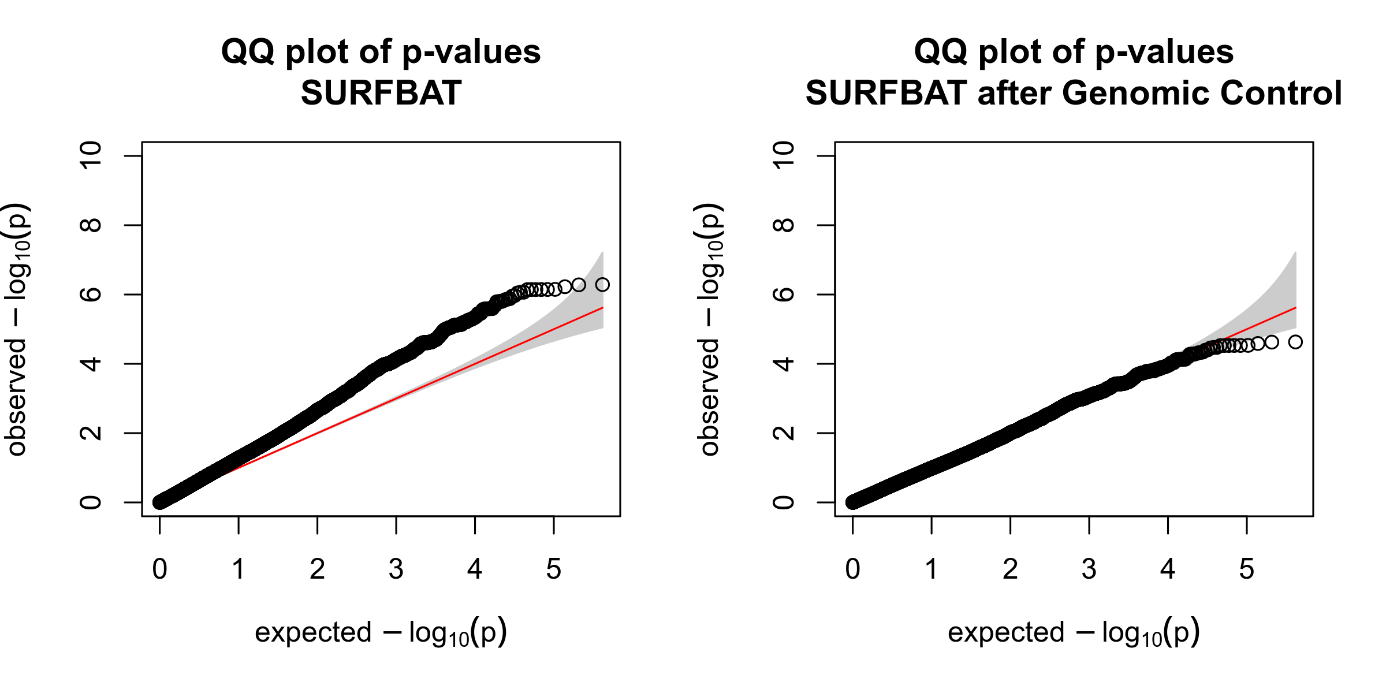

Supplementary Figure 2b: Using the data from chromosome 1 corresponding to Supplementary Figure 1a, the resulting inflation in the SURFBAT test-statistic is shown (left). Applying Genomic Control (right) successfully removes this inflation.

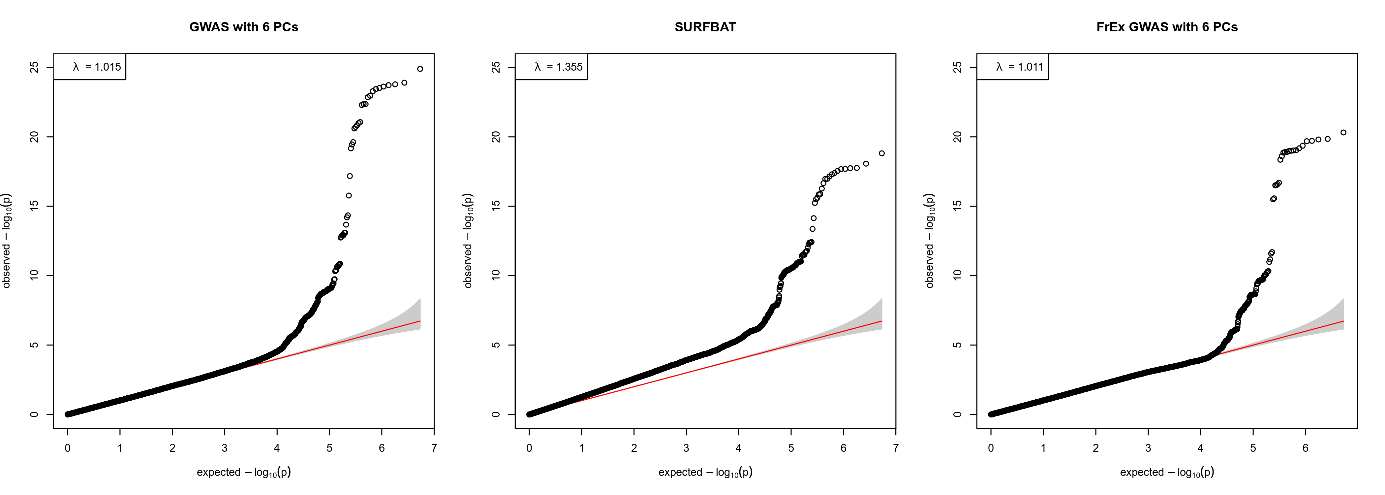

Supplementary Figure 3:

QQ-plots corresponding to Figure 2 in the main text before genomic control has been applied.
